# Molecular characterization of the viral structural gene of the first dengue virus type 1 outbreak in Xishuangbanna, a border area of China, Burma and Laos

**DOI:** 10.1101/423152

**Authors:** Yao Lin, Dehong Ma, Songjiao Wen, Fen Zeng, Shan Hong, Lihua Li, Xiaoman Li, Xiaodan Wang, Zhiqiang Ma, Yue Pan, Junying Chen, Juemin Xi, Lijuan Qiu, Xiyun Shan, Qiangming Sun

**Author notes:** Yao Lin and Dehong Ma contributed equally to this article. **Correspondence:** Qiangming Sun, Xiyun, Shan.

## Abstract

In the context of recent arbovirus epidemics, dengue fever is becoming a greater concern around the world. In August 2017, Xishuangbanna, which is a border city of China, Burma and Laos, had its first major dengue outbreak. A total of 156 serum samples from febrile patients were collected; 97 DENV positive serum samples were screened out, and viral RNAs were successfully and directly extracted, including 77 cases from China and 20 cases from Myanmar. Phylogenetic analysis revealed that all of the strains were classified as DENV-1. There are eight epidemic dengue strains from Myanmar and 74 from Jinghong, Xishuangbanna, that were classified as cluster 1, which are the most similar to the strain of China Guangzhou 2011. There are three epidemic strains from Xishuangbanna Mengla that were classified as cluster 2, which have the closest relationship to the strain of China Hubei 2014. However, there are 12 epidemic strains from Myanmar that were classified as cluster 3, which have the closest relationship to the strain of Laos from 2008, which shows that there is a recycling epidemic trend of DENV in China. There were 236 mutations in the base, which caused 31 nonsynonymous mutations in the DENV structural protein C/prM/E genes when the strain of Xishuangbanna and Myanmar were compared with the DENV-1SS. There is no clear homologous recombination signal among these stains. Homology modeling possibly predicted a three-dimensional structure of the structural protein of these strains and revealed that they had the same three-dimensional structure and all had five predicted protein binding sites, but there are differences in binding site 434 (DENV-1SS: Thr434, DV-Jinghong: Ser434, DV-Myanmar: Ser434, DV-Mengla: Ser434). The results of the molecular clock phylogenetic and demographic reconstruction analysis show that DENV-1 became highly diversified in 1972 followed by a slightly decreased period until 2017. In conclusion, our study lays the foundation for studying the global evolution and prevalence of DENV.

**Author Summary:** Dengue fever (DF) is a mosquito-borne illness caused by a flavivirus. Human infections with Dengue virus (DENV) could cause fever, cutaneous rash and malaise. Xishuangbanna, which is located in the southwestern Yunnan Province and is a border city with China, Burma and Laos, was reported to have outbreak of DENV in 2013 and 2015 with different types. However, there was a large outburst of dengue in May 2017. To understand the genetic characterization, potential source and evolution of the virus, 156 serum samples were analyzed. We focused on: (i) Phylogenetic analysis of the structural protein genes sequences; (ii) Mutation, recombination analysis and predicted protein binding sites of the structural protein genes; (iii) Molecular clock and demographic reconstruction of global dengue virus serotype 1(DENV-1). Our results indicated that this is the first outbreak of DENV-1 in Xishuangbanna, dengue epidemic strains on the Burma border of China show diversification, we found a virulence site changed from I to T(amino acid position: 440), which may lead to weakened virulence of the epidemic strains. We found that the evolution of DENV-1 is dominated by regional evolution. What’s more, DENV-1 became highly diversified in 1972 followed by a slightly decreased period until 2017.

## Introduction

Dengue virus (DENV) is one of the most important human arboviruses and is primarily transmitted by mosquitoes *Aedes albopictus* and *Ae. aegypti* [1]. Dengue virus causes dengue fever (DF), which is a major global public health problem that is endemic in the world. Dengue virus encodes three structural proteins (C/prM/E) and seven non-structural (NS1, NS2A, NS2B, NS3, NS4A, NS4B, and NS5) proteins. There are four serotypes (DENV-1, DENV-2, DENV-3 and DENV-4) that are prevalent throughout the tropical and subtropical regions of the world [2,3,4]. With a 30-fold increase in global incidence over the past 50 years, it is estimated that 390 million dengue infections occur annually worldwide [5,6].

The first outbreak of dengue in mainland China occurred in Foshan of the Guangdong Province in 1978 [5,7,8]. Since that outbreak, disease outbreaks have been continually recorded in the Hainan, Fujian, Guangxi and Zhejiang Provinces [6,9]. Yunnan province in southwestern China and borders the countries of Southeast Asia, where dengue fever (DF) is common. The first outbreak in Yunnan Province was reported in 2008 with 56 confirmed cases. Since that outbreak, epidemics have been regularly reported in Yunnan [10]. Xishuangbanna, which is located in the southwestern Yunnan Province and is a border city with China, Burma and Laos, was reported in 2013 to have the first outbreak with 1319 confirmed DENV-3 cases [11,12,13]. Later, 1132 confirmed DENV-2 cases were reported in Xishuangbanna in 2015 [14]. However, there was a large outburst of dengue in May 2017. The epidemic continued until the middle of November at which point 1348 cases were confirmed. At the same time, DENV also broke out in Myanmar, which is contiguous with Xishuangbanna. Accordingly, the information of origin and molecular epidemiology involved with DENV deserves to be studied.

In this study, we report the results of the detection and serotyping of the DENV that caused the outbreak. Phylogenetic and molecular clock analyses based on structural gene sequences of the virus were also performed to understand the genetic characterization, potential source and evolution of the virus.

## Materials and Methods

### Ethical statement

All of the participants were informed of the study aims, and written informed consent was received from each patient before sample collection. The study protocol was approved by the Institutional Ethics Committee (Institute of Medical Biology, Chinese Academy of Medical Sciences, and Peking Union Medical College) and was in accordance with the Declaration of Helsinki for Human Research of 1974 (last modified in 2000). There are 146 participants were adults, 10 participants were minors. 10 minors’ parents provided consent on behalf of all child participants.

### Dengue virus identification and viral RNA Extraction

Serum samples of dengue fever patients were collected from blood and were followed by viral RNA extraction. A rapid dengue fever NS1 Test Kit (Blue Cross, Beijing, China) was used to detect the DENV NS1 antigen. The QIAamp Viral RNA Mini Kit (Qiagen, Hilden, Germany) was used for viral RNA extraction, 150 μl of each serum sample was used and following the manufacturer’s instructions, and the RNA was finally saved at −80 °C.

### Reverse transcriptase-polymerase chain reaction (RT-PCR) and sequencing of the C/prM/E gene

For the genotyping and phylogenetic analysis, the Prime Script^TM^™ 1^st^ Strand/cDNA Synthesis Kit (TaKaRa Co., Ltd. Dalian, China) was used followed by the manufacturer’s instructions to synthesize the cDNA. Two oligo nucleotide primer pairs were synthesized through Primer-BLAST on the NCBI web (http://www.ncbi.nlm.nih.gov/tools/primer-blast/). A total of 2322 bp gene sequences of the DENV structural proteins C/prM/E were obtained with a reference to the standard strain DEN1SS (GenBank ID: DQ672562) to amplify the overlapping fragments between base sites 98–2419.

The primer sequences were as follows: C/prM-Forward, 5′ - agttgttagtctacgtggac -3′ and C/prM-Reverse, 5′-agagttcaatgtccagtgtt-3′; E-Forward, 5′-gattcacggtgatagccctt -3′ and E-Reverse, 5′ - gctctgtccaggtgtggact -3′. All of the primers were synthesized and purified by an outside vendor (TsingKe Biotech Co, Ltd. Beijing, China).

### Virus isolation

Virus isolation was performed by inoculation into the C6/36 *Aedes albopictus* cell line (ATCC CRL-1660) to construct the dengue viral seeds library.

### Sequences and phylogenetic analysis

Sequences of the structural protein genes (C/prM/E) were aligned by the ClustalX program and compared with the reference strains, which were collected from GenBank under the following accession numbers 43: Standard strain (DENV-1SS:DQ672562, DENV-2SS:M29095, DENV-3SS:M93130, and DENV-4SS:AF326573); China (JQ048541, KU365900, KP772252, DQ193572, FJ176780, AB608788, KU094071, and KT827374); Malaysia (KX452065 and KX452068); Singapore (GU370049, KJ806945, GQ357692, M87512, JN544411, and AB204803); South Korea (KP406802); Indonesia (KC762646); Japan (AB178040); Laos (KY849701, KC172832, KC172829, KC172835, and KC172834); Thailand (HM469966); Myanmar (AY726554, AY726553, and JF459993); Cambodia (HQ624984); Vietnam (JQ045660); USA (DQ672564, EU848545, KT279761, FJ390379, and AF180817); Brazil (AF226685); India (KF289072); Nauru Island (U88535); and French Polynesia (JQ915076). Next, phylogenetic analysis was performed using Molecular Evolutionary Genetics Analysis (MEGA) software version 6.0 through the Maximum Likelihood phylogeny test with a bootstrap of 1000 replications. Afterwards, a Bayesian skyline plot (BSP) analysis in BEAST software was used to build an evolutionary tree to identify the results of the ML tree.

### Molecular characterization

These sequences of the structural proteins of the 2017 Yunnan and Myanmar DENV epidemic strains were deposited in the NCBI GenBank database (http://www.ncbi.nim.nih.gov/GenBank/index.html): MG696640-MG696656 and MG523189-MG523268. Next, the nucleotide sequence substitutions and translated amino acid sequence mutations were analyzed with BioEdit and the Sequence Manipulation Suite (SMS, http://www.bio-soft.net/sms/index.html). SimPlot was used to predict the homologous recombination in the structural proteins of the DEN1SS and the other 2017 epidemic strains including MG523256, MG523190, and MG696653.

Phyre^2^ (http://www.sbg.bio.ic.ac.uk/phyre2/html) was used to predict ligand binding sites and amino acid variation in the three-dimensional structure between structural proteins of DEN1SS and 2017 Xishuangbanna and Myanmar epidemic strains.

### Molecular clock and demographic reconstruction

A number of available full-length DENV genome sequences were downloaded from GenBank under the following accession numbers: China (JQ048541, KU365900, KP772252, DQ193572, FJ176780, AB608788, KU094071, and KT827374); Malaysia (KX452065 and KX452068); Singapore (GU370049, KJ806945, GQ357692, M87512, JN544411, and AB204803); South Korea (KP406802); Indonesia (KC762646); Japan (AB178040); Laos (KC172829, KC172835, and KC172834); Thailand (HM469966); Myanmar (AY726554 and AY726553); Cambodia (HQ624984); Vietnam (JQ045660); USA (DQ672562, EU848545, KT279761, FJ390379, and AF180817); Brazil (AF226685); India (KF289072); Nauru Island (U88535); French Polynesia (JQ915076); along with MG523190, MG523256 and MG696653 in our study, and the alignment was performed to obtain the structural protein region. Phylogenetic analysis and demographic reconstruction were collectively estimated using Bayesian Evolutionary Analysis by BEAST and TreeAnnotator and using strict and uncorrelated lognormal relaxed models, respectively. For population dynamics, the Bayesian skyride model was used. One million steps were run, and the first 10% were removed as burn-in.

## Results

### Geographic analysis between Xishuangbanna and Myanmar and the study design

A geographical analysis was performed among countries of Yunnan China, Guangzhou China and bordering Southeast Asian countries, including Myanmar, Laos and Vietnam. The results from the geographical analysis show that Yunnan has become a central DENV epidemic area and Xishuangbanna is adjacent to South East Asian countries. The samples in our study were collected from two counties in Xishuangbanna, namely, Mengla and Jinghong (Fig. 1). From August to November 2017, dengue fever broke out in Xishuangbanna and Myanmar with a total of 1179 dengue patients at Xishuangbanna Dai Autonomous Prefecture People’s Hospital. A total of 156 serum samples from inpatients were collected from this Hospital and Menghai Daluo Health-center, and all of these 156 cases were confirmed to be NS1 positive by the Colloidal Gold test. Among them, 97 DENV positive serum samples were screened from patients whose fever courses were shorter than 5 days. Among these strains, a total of six virus strains led to cytopathic effects after amplification in C6/36 cells for 6 days by continuous passage. In addition, 97 viral RNAs were successfully and directly extracted from these serum samples following gene sequencing of the DENV structural protein C/prM/E genes. Analysis of the phylogenetic, mutation, homologous recombination signal and possible three-dimensional structure and molecular clock analyses were then conducted successively as shown in Fig. 2.

**Figure 1.**
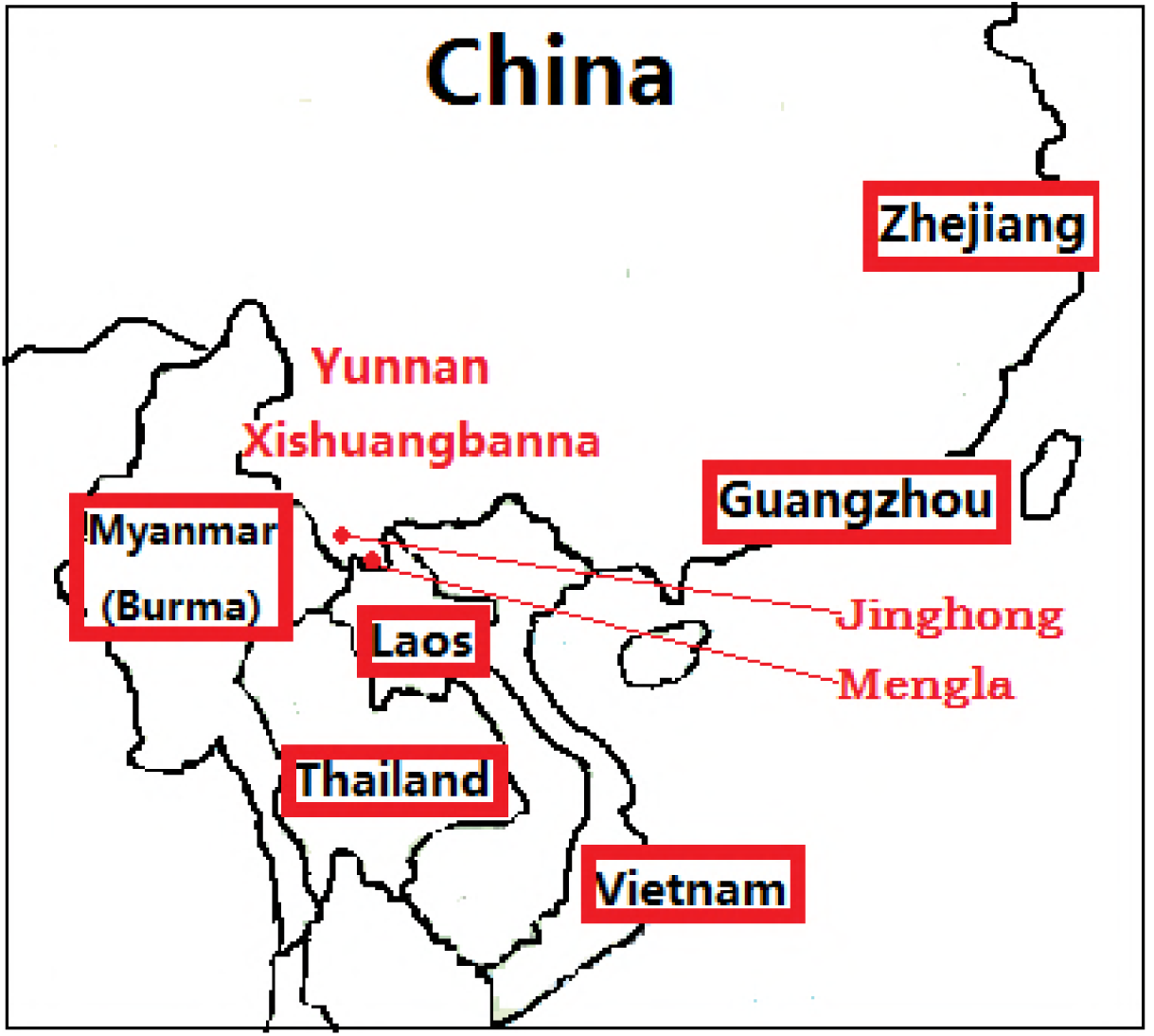
Geographic relationships between Xishuangbanna and other Asian dengue outbreak countries and areas in 2017. The red diamond was used to mark some dengue outbreaks area and the red block represents the other dengue epidemic areas in China and around Asia. Paint software of Microsoft Windows 6.1 was used to plot this graph. The graph showed that Yunnan province was geographically a connecting area among them, which led to possible dengue epidemic in Xishuangbanna.

**Figure 2.**
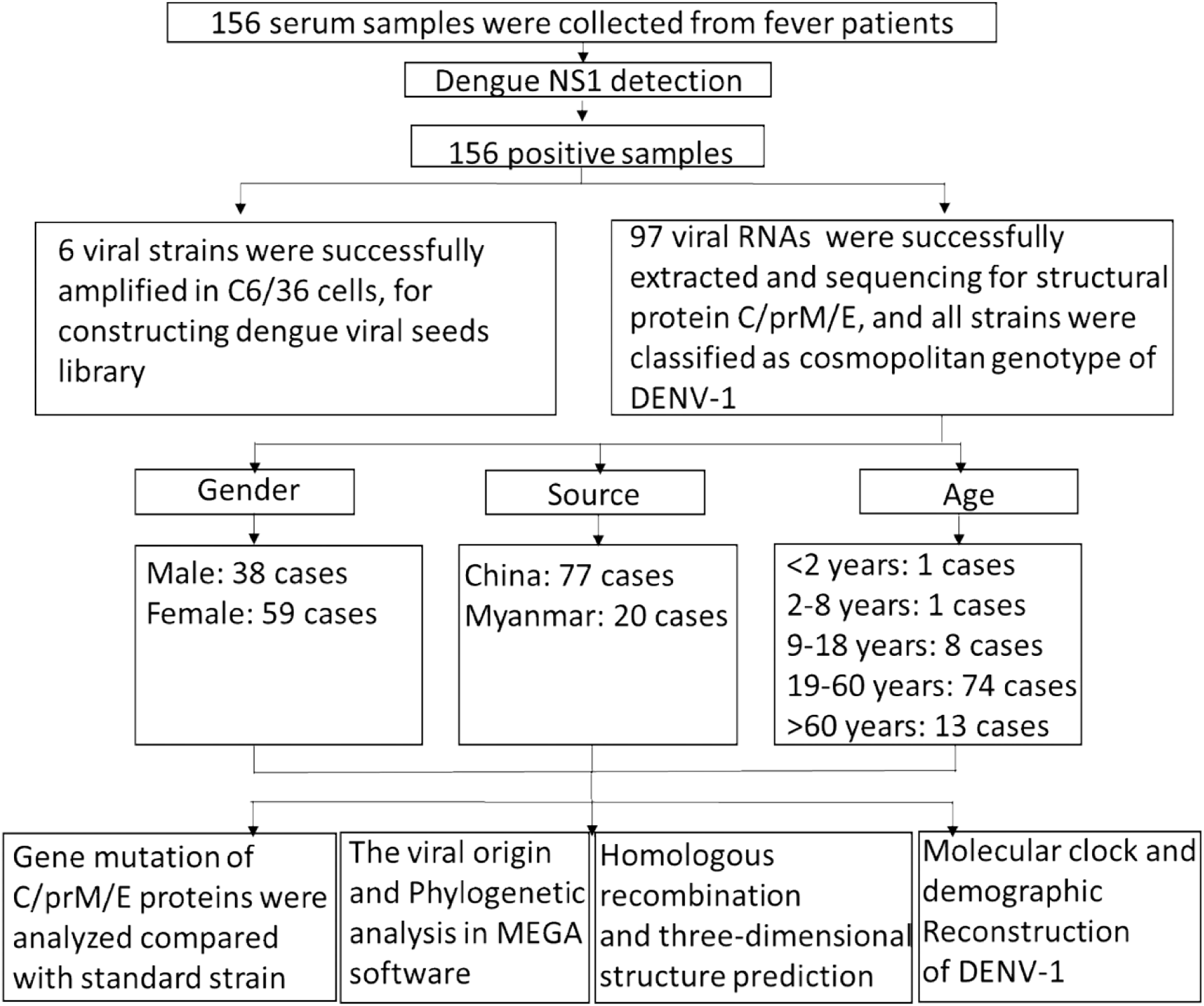
The study design and the following analysis. 156 inpatients who was diagnosed with dengue fever were recruited in our study; among them, all cases were first identified as dengue NS1 positive. Of these, serum samples were collected for virus amplification and viral RNAs extraction followed by Phylogenetic analysis to characterize the origin and prevalence of this DENV outbreak.

### Phylogenetic analysis

All of the structural protein genes sequences, which could be successfully extracted viral RNA, were aligned to conduct the phylogenetic analysis. In addition to these 97 structural protein gene sequences, the relevant reference sequences were also obtained from the GenBank database. Typical Chinese strains, other epidemic strains from neighboring countries and standard strains of four serotypes were included. The results of the phylogenetic analysis show that all 97 strains of dengue virus were serotype 1 DENV and can be divided into three clusters in the ML tree. There are eight epidemic dengue strains from Myanmar, and 74 from Jinghong Xishuangbanna were classified as cluster 1, which are most similar to the strain of China Guangzhou 2011 (JQ048541). There are three epidemic strains from Xishuangbanna Mengla that were classified as cluster 2, which have the closest relationship to the China Hubei 2014 (KP772252) strain. However, there are 12 epidemic strains from Myanmar that were classified as cluster 3, which have the closest relationship to the strain of Laos 2008 (KC172834) (Fig. 3). The result indicates that dengue fever is transmitted by the close proximity of the countries. In addition, dengue fever has become a disease circulating in the interior of China.

**Figure 3.**
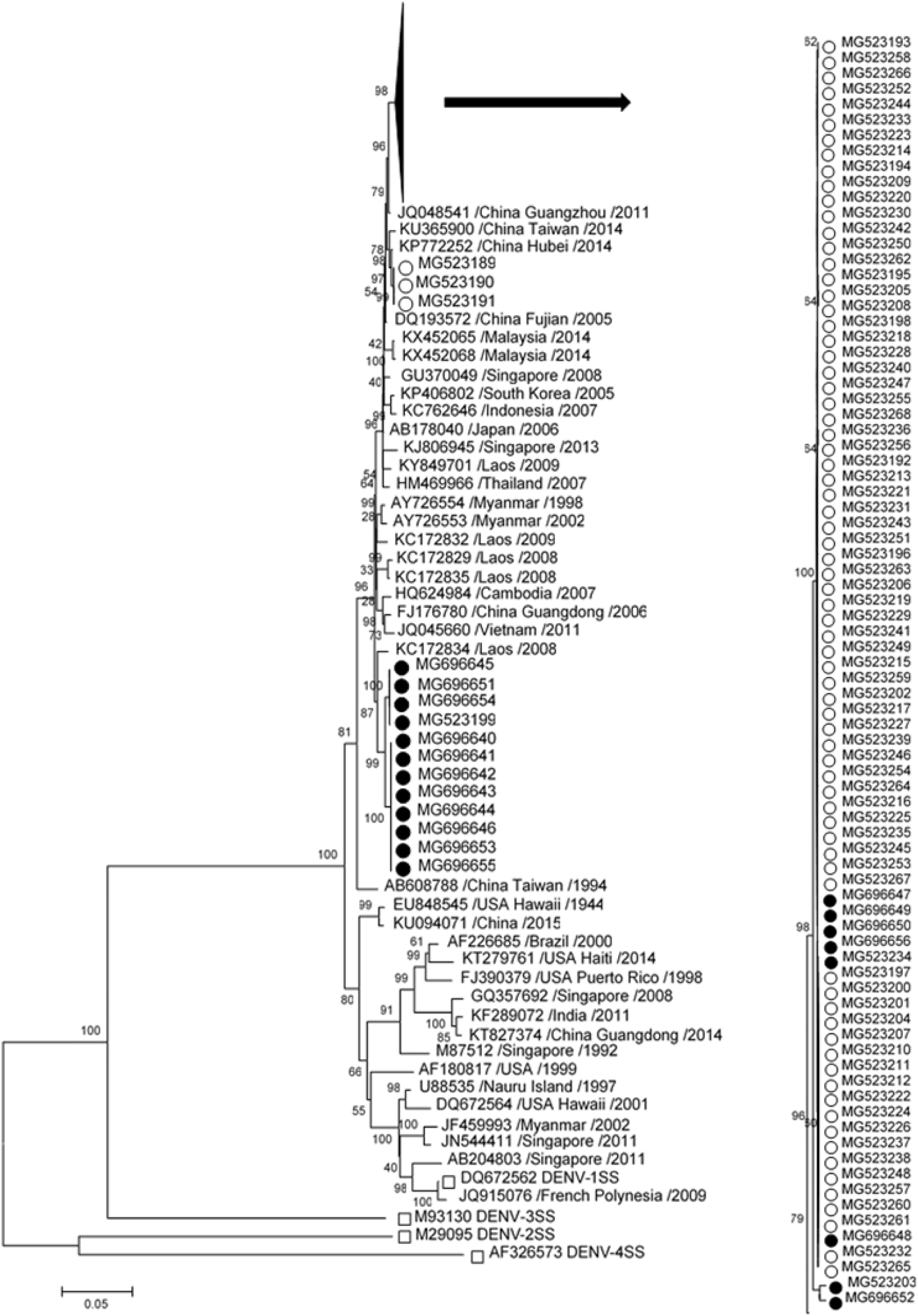
Phylogenetic tree of DENV-1 epidemic strains in Xishuangbanna, Yunnan, China and Myanmar, 2017. The phylogenetic tree was constructed using the Maximum Likelyhood phylogeny test. Bootstrap values were set for 1000 repetitions. (● denotes DENV strains from Myanmar 2017, ○ denotes DENV strains from Xishuangbanna, Yunnan, China in 2017, □ denotes the standard strains of four dengue serotypes. The sequences of the reference strains were derived from the NCBI GenBank database (www.ncbi.nlm.nih.gov).

### Mutation and recombination analysis

First, the 97 sequences of structural protein coding regions were found to be most like DENV-1SS instead of the other three standard strains by blast and sequence similarity comparison. The nonsynonymous mutation of the DV-Jinghong, DV-Myanmar, DV-Mengla strains were tested and compared to the DENV-1SS reference strain. The result shows that there were 236 mutations in the base causing 31 nonsynonymous mutations in the DENV structural protein C/prM/E genes. These differences in molecular characteristics of DV-Jinghong, DV-Myanmar, and DV-Mengla strains are considered the point of penetration for the study of the dengue outbreak in Xishuangbanna and Myanmar (Fig. 4a). The possible homologous recombination of C/prM/E proteins of dengue virus 2017 were then predicted and compared with DENV-1SS. There are no obvious homologous recombination signals among these strains (Fig. 4b).

**Figure 4.**
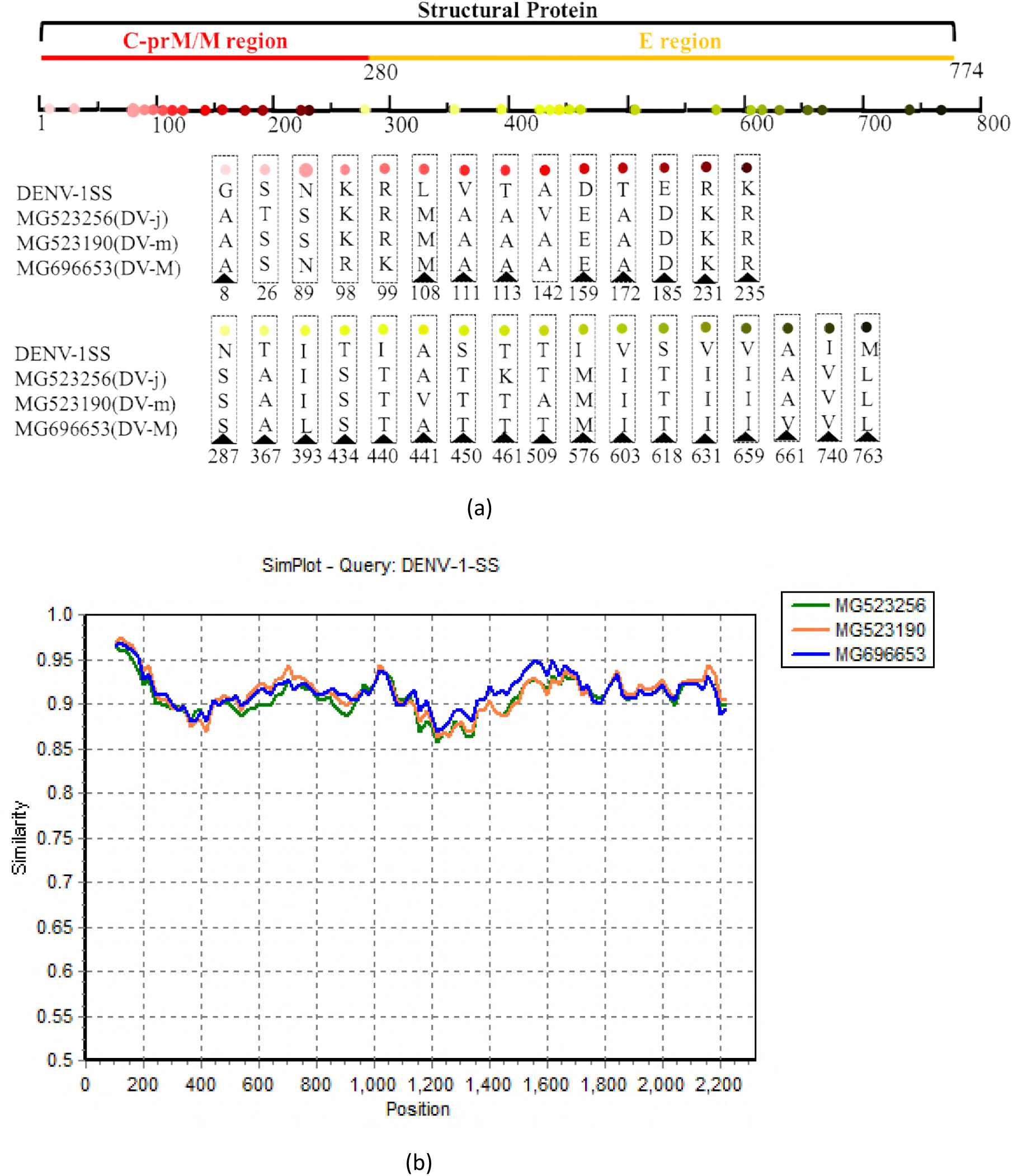
Molecular character analysis of the DV-Jinghong,DV-Myanmar, DV-Mengla strains 2017 comparing with DENV-1 standard strain. (**a**) amino acid mutations of the DV-Jinghong, DV-Myanmar, DV-Mengla strain compared with the DEN1SS. DV-j, DV-m, DV-M indicated DV-Jinghong, DV-Mengla, DV-Myanmar respectively. ▲ Indicates that the mutation existed in all sample strains. (**b**) Homologous recombination prediction of the structural proteins for DEN1SS and DV-Jinghong, DV-Myanmar, DV-Mengla. MG523256 represents the DV-Jinghong, MG523190 represents the DV-Mengla, MG696653 represents the DV-Myanmar. No obvious homologous recombination signal among these stains.

### Possible three-dimensional structure of the structural protein C/prM/E genes

The possible three-dimensional structure of the structural protein of DV-Jinghong, DV-Myanmar, and DV-Mengla were later predicted and compared with DEN1SS. Homology modeling revealed that four strains had the same three-dimensional structure, and all five had predicted protein binding sites, which were DENV-1SS (His428, Gln429, Glu433, Thr434, and Thr435), DV-Jinghong (His428, Gln429, Glu433, Ser434, and Thr435), DV-Myanmar (His428, Gln429, Glu433, Ser434, and Thr435), and DV-Mengla (His428, Gln429, Glu433, Ser434, and Thr435). We found one of the binding sites is different, i.e., 434 (Fig. 5).

**Figure 5.**
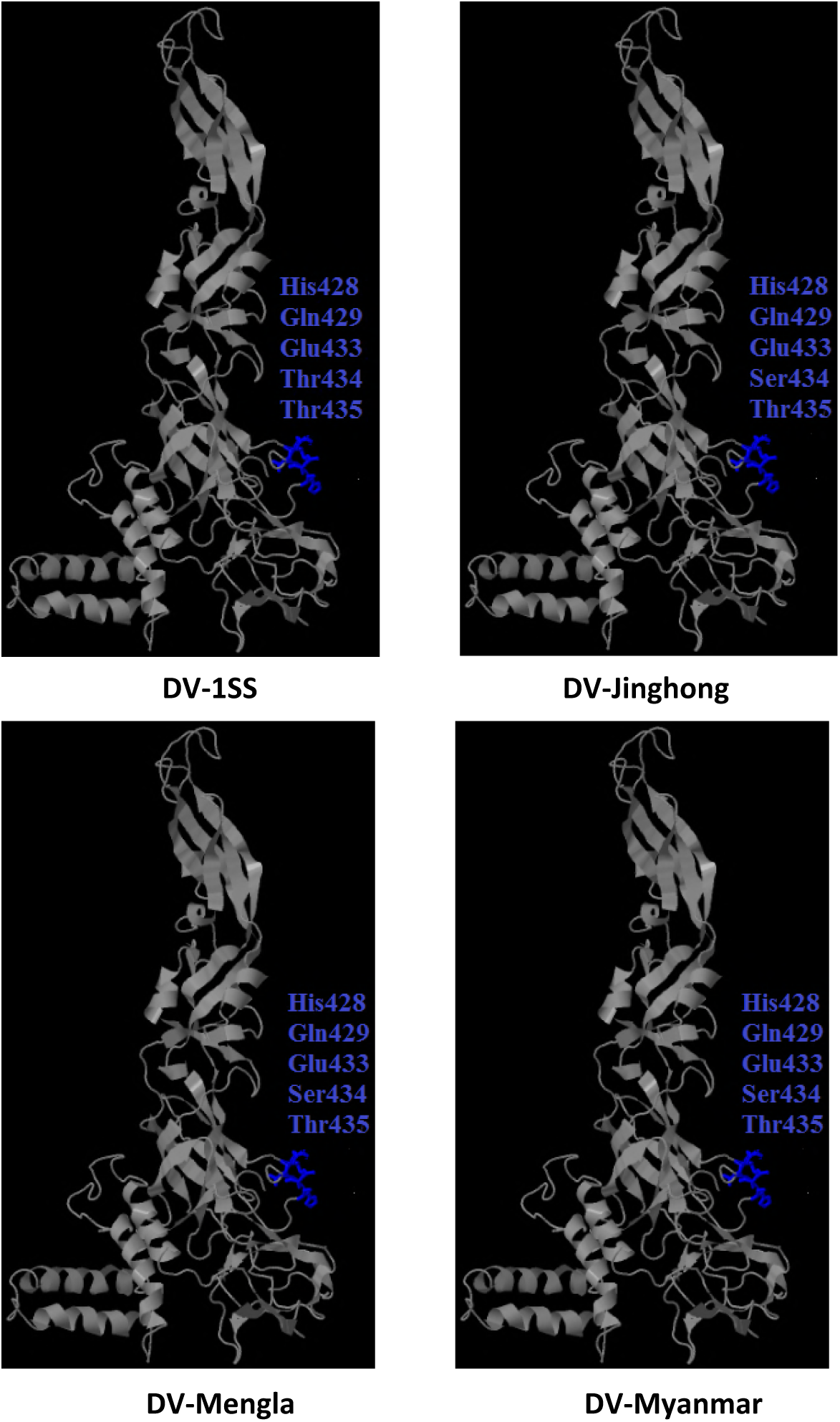
Possible three-dimensional structure of structural protein C/prM/E genes for DEN1SS and DV-Jinghong, DV-Mengla, DV-Myanmar. DV-1SS indicates DENV-1 standard strain, DV-Jinghong, DV-Mengla, DV-Myanmar indicated that the dengue epidemic occurred in Xishuangbanna Jinghong, Xishuangbanna Mengla, Myanma, respectively. Blue indicates predicted protein binding sites. There are 5 possible protein binding sites in the structural protein region of the four strains.

### Molecular clock and demographic reconstruction

The results of phylogenetic analysis show that all strains of dengue virus serotype 1 share a common ancestor and with evolution can be divided into two categories in 1867 (Figure 6A) (in cluster 1), and most of them are the dengue that broke out in America. However, in cluster 2, most of these strains are DENV-1 that broke out in China and the countries around China, such as Myanmar and Laos. This finding indicates that the evolution of DENV-1 is dominated by regional evolution. In cluster 2, DENV-1 became highly diversified in 1972 (Figure 6A), which was supported by the coexistence of several minor clusters, and three typical viruses (DV-Jinghong, DV-Myanmar, and DV-Mengla) in our study have common ancestry at this time. The evidence for increasing genetic diversity was also supported by the demographic reconstruction analysis (Figure 6B). Since 1961, the genetic diversity of DENV-1 has gradually increased and reached its peak in approximately 1972 following a slightly decreasing period until 2017. During this period and although the analysis based on the currently available data suggests that the genetic diversity decreased slightly after 1972 (Figure 6B), we consider that this could be caused by the extremely low sampling density in cluster 1.

**Figure 6.**
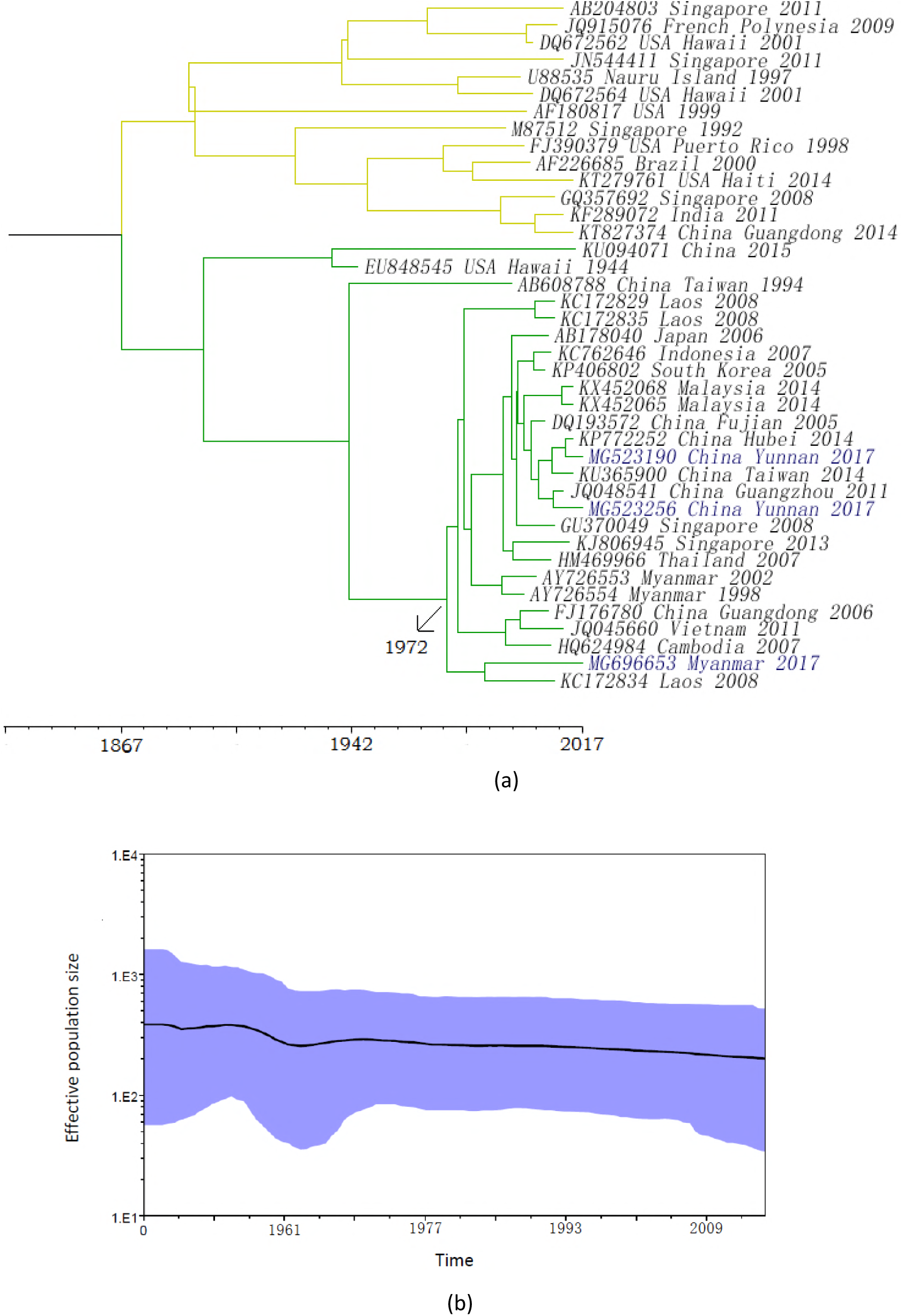
Phylogenetic analysis and demographic reconstruction of DENV-1 genome sequences. Phylogenetic trees of DENV-1 structural genome sequences from all over the world since 1944. (a) BEAST using a strict molecular clock. viruses in our study are labeled in blue, MG523190 represents the DV-Mengla, MG523256 represents the DV-Jinghong, MG696653 represents the DV-Myanmar. Branch 1 is labeled in yellow, and branch 2 in green. (b) The demographic reconstruction of the DENV-1 structural genome sequences since 1944. The blue interval represents the HPD of 95%. This was estimated using the uncorrelated lognormal relaxed molecular clock and the GMRF Bayesian skyride model.

## Discussion

In China during 1978–1991, the epidemic regions were primarily in Guangdong and Hainan. After 1990, the endemic range of dengue fever in China expanded geographically, with new cases being reported in Guangxi, Fujian, Zhejiang, Jiangsu, Yunnan, Henan, Hubei, and Beijing in recent years.

According to the National Notifiable Disease Surveillance System, from 2002 to 2016, the incidence ranged from 0.01 to 3.46 per 100,000 people with a total of seven deaths reported. In 2014, there were more than 44,000 reported cases of dengue fever, which stands out as the highest-ever number of dengue infections per year in the historical records [15,16]. Yunnan Province is located in southwestern China and neighbors the southeast Asian countries, which are all dengue-endemic areas. Research indicates that imported DF from patients from Laos and Myanmar was the primary cause of the DF epidemic in Yunnan Province [17,18,19], whereas Chinese encouragement of implementing economic development policy with Southeast Asian will result in greater contacts between Yunnan and southeast Asian countries. This finding indicates that the dengue epidemic might become increasingly common in this Chinese border area.

Xishuangbanna is located in southwestern Yunnan Province, which is a border area of China, Burma and Laos. In 2013 and 2015, DENV-3 and DENV-2 outbreaks were reported in Xishuangbanna with 1319 and 1132 infections, respectively [20,21,22]. In 2017, the first DENV-1 outbreak in the history of Xishuangbanna occurred. It is infrequent for an outbreak to occur at the scale of the DENV-1 epidemic at these locations. A total of 156 serum samples were collected from febrile patients at Xishuangbanna Dai Autonomous Prefecture People’s Hospital and Menghai Daluo Health Center, and all of the cases were confirmed to be dengue NS1 positive by the most authoritative local hospital on DENV treatment. Because approximately 72% of the 156 serum samples were collected in the early stage of the disease and with a febrile course shorter than 3 days, DENV RNA was successfully extracted from 97 serum samples. However, the success rate of virus isolation from the serum of these patients is extremely low, and only six viral strains were successfully amplified in the C6/36 cells for construction of the dengue viral seeds library. DENV infected cells can show a visible cytopathic effect (CPE) by serial-propagation. It is worth noting that the DENV outbreak broke out in different years and varied with different types in Xishuangbanna. This phenomenon warrants further study, especially to examine the influence factor among mosquitoes, the environment and tourism in Xishuangbanna.

The phylogenetic analysis was conducted based on the structural protein genes of the epidemic strain, and several epidemic strains from China and Southeast Asian countries were used as references. Previous studies report that the imported cases in China (2013-2016) were mainly from Asian countries, such as Burma and African countries, which also varied with the type of infectious disease [23,24,25]. In addition, epidemiological and molecular characteristics of emergent dengue virus in Yunnan Province (2013-2015) indicated that imported DF from patients of Laos and Myanmar were the primary cause of the DF epidemic in the Yunnan Province [12]. The results of the phylogenetic analysis show that all 97 strains of dengue virus were serotype 1 DENV and can be divided into three clusters in the ML tree. The outbreak of the dengue epidemic caused by insect vectors and frequent travel is beyond a doubt. Evolutionary analysis is used to locate the regions of a protein that are important for its function or structure. The rate of evolution is generally constant for a given family of homologous sequences [26]. In our study, these results will help us better understand dengue’s evolution and the prevalent trend in Xishuangbanna. Changes in the hydrophobicity of amino acids can cause major changes in antigenicity and may affect the virulence of the virus [27,28]. Rey et al. determined the three-dimensional structure of the E protein of the tick-borne encephalitis (TBE) virus, and this structure can represent the three-dimensional structure of the E protein of the flavivirus [29]. In this structure, the E protein consists of three domains: I region: the central region carrying the antigenic site, II: the dimer fusion region, and III: likely related to the binding of cellular receptors. The E155 amino acid residue of the DEN-4 virus is located in the domain I region. Kawano et al. used dengue type 4 high-virulence and low-neuroviral viruses to construct an internal chimeric virus and perform point mutation analysis. After finding that amino acid residue E155 was converted from T to I, one of the two potentially conserved N-linked glycosylation sites of the E protein (Asn-Ile-Thr155) changes, which changes the antigenic site of the I region and increases the virulence of the virus [30]. In our study, the E155 locus corresponds to the DENV-1 structural protein (amino acid position: 440). Strains of dengue fever in Xishuangbanna and Myanmar in 2017 were compared with DENV-1SS. The virulence site changed from I to T, which may lead to weakened virulence of the epidemic strains. This is also consistent with the very rare phenomenon in the outbreak of severe dengue patients.

Gene recombination is an important mechanism for the evolution of the virus. Through gene recombination, a virus can produce great genetic variation, and its speed is clearly higher than that caused only by mutation [31,32,33]. To reveal the role of genetic mechanisms, such as gene recombination in genetic evolution, the structural protein of DENV-1SS DV-Jinghong, DV-Mengla, and DV-Myanmar were analyzed by homologous recombination software SimPlot. The results show that there are no recombinant signals in the four strains of DENV. Homology modeling prediction of a possible three-dimensional structure of the structural protein of these strains revealed that they had the same three-dimensional structure, and all had five predicted protein binding sites, but there are differences in binding site 434 (DENV-1SS: Thr434; DV-Jinghong: Ser434; DV-Myanmar: Ser434; and DV-Mengla: Ser434). The results of the phylogenetic and demographic reconstruction analysis show that DENV-1 became highly diversified in 1972 followed by a slightly decreasing period until 2017. In conclusion, our study lays the foundation for studying the global evolution and prevalence of DENV.

In 2013, 2015 and 2017, interestingly, there have been extensive outbreaks of dengue fever in Xishuangbanna, which is a border area of China, Burma and Laos with different serotypes. In 2017, a dengue outbreak expanded in China and included Zhejiang, Shandong, Yunnan, Guangdong, Xianggang, Taiwan, Macao and Southeast Asian countries close to Xishuangbanna. The tracing and propagation of epidemic viruses is highly complicated. Our study should serve as a reference to follow-up studies of molecular epidemiology, virulence, infection, and pathogenicity of DENV.

## Acknowledgments

We would like to thank Dr. Penghua Wang of Department of Microbiology and Immunology, School of Medicine, New York Medical College for proofreading and reviewing our manuscript.

